# Schizophrenia-associated genomic copy number variants and subcortical brain volumes in the UK Biobank

**DOI:** 10.1101/374678

**Authors:** Anthony Warland, Kimberley M Kendall, Elliott Rees, George Kirov, Xavier Caseras

**Author notes:** Corresponding Author: Dr Xavier Caseras, MRC Centre for Neuropsychiatric Genetics and Genomics, Cardiff University, Hadyn Ellis Building, Maindy Road, Cardiff, CF24 4HQ, UK.

## Abstract

Schizophrenia is a highly heritable disorder for which anatomical brain alterations have been repeatedly reported in clinical samples. Unaffected at-risk groups have also been studied in an attempt to identify brain changes that do not reflect reverse causation or treatment effects. However, no robust associations have been observed between neuroanatomical phenotypes and known genetic risk factors for schizophrenia. We tested subcortical brain volume differences between 49 unaffected participants carrying at least one of the 12 copy number variants associated with schizophrenia in UK Biobank and 9,063 individuals who did not carry any of the 93 copy number variants reported to be pathogenic. Our results show that CNV carriers have reduced volume in some of the subcortical structures previously shown to be reduced in schizophrenia. Moreover, these associations were partially accounted for by the association between pathogenic copy number variants and cognitive impairment, which is one of the features of schizophrenia.

## Introduction

Schizophrenia (SZ) is a severe psychiatric disorder with a profound impact on affected individuals, their families and society. Patients with SZ have an average life expectancy 10-20 years shorter than the general population, around 20% experience chronic psychotic symptoms, 50% long-term psychiatric problems and 80-90% unemployment^1–4^. Despite research efforts, our ability to treat SZ remains limited and no new therapeutic targets of proven efficacy have been developed for decades. Clozapine, discovered almost six decades ago, still stands as the most effective treatment for SZ^5^. To a large extent, this lack of progress is due to our poor understanding of the neurobiological underpinnings of the disorder; although recent developments in brain imaging have opened new opportunities to advance on this knowledge. Multiple case-control studies have consistently shown the existence of anatomical abnormalities in the brains of individuals with SZ, although limitations in statistical power and sample representativeness have resulted in frequent discrepancies with regards to a more detailed description of these abnormalities. Overcoming these limitations, ENIGMA – a large international consortia effort^6^ - recently showed reduction in the volume of the hippocampus, thalamus, amygdala and accumbens, but increased volume in lateral ventricles and pallidum, in individuals with SZ relative to controls^7^; results largely replicated by a parallel Japanese consortium (COCORO)^8^. However, it remains possible that findings from case-controls studies reflect reverse causation and/or treatment effects rather than causal neurobiological mechanisms.

Since SZ is a highly heritable disorder^9^, it would be expected that any potential marker of pathogenic mechanisms would also be detectable in participants at high genetic risk who are unaffected by SZ. Important advantages of this approach lie in its ability to access large samples of participants and avoid the possibility of reverse causation or treatment effects. A recent study probed the association between common genetic risk for SZ, i.e. polygenic score based on the latest GWAS data available^10^, and subcortical brain volumes in a sample of over 11,000 participants^11^, but found no significant associations, calling into question whether any of the above ENIGMA findings were driven by genetic risk. Although common risk variants collectively account for a greater proportion of the genetic liability to SZ than known rare risk variants, such as copy number variation (CNV), individually the latter show much higher penetrance for the disorder (ranging between 2-18%)^12^. Therefore, rare risk CNVs might be better suited to capture some of the brain anatomical variability associated with SZ. Following the above study^11^, we aimed to investigate the association between genetic risk for SZ, defined as carrying at least one CNV known to increase risk for SZ, and volume of subcortical brain structures in a large sample of ~10,000 unaffected participants from the UK Biobank project. We hypothesised that individuals carrying SZ-associated CNVs (SZ-CNVs) will have smaller volume in hippocampus, thalamus, amygdala and accumbens, but larger in pallidum. Volume of lateral ventricles is not currently an available measure from UK Biobank participants; therefore, we use overall ventricular volume as a proxy. We based our selection of SZ-CNVs on previously published work^13–16^, which found 12 specific CNVs to be significantly associated with SZ after correction for multiple testing. Since cognitive impairment is a core feature of SZ and previous research has shown genetic overlap between IQ and SZ^17^, we also aimed to examine whether a previously reported association between SZ-CNVs and cognitive performance^18^ was mediated by subcortical volume alterations. Of the cognitive measures available from scanned participants in UK Biobank, we selected fluid intelligence (FI) score due to its strong heritability and robustness against influences from education and training^19^.

## Materials and Methods

### Participants

This study used a subsample of participants from UK Biobank (www.ukbiobank.ac.uk). All subjects provided informed consent to participate in UK Biobank and agreed to follow-up assessments. Ethical approval was granted by the North West Multi-Centre Ethics committee. Data were released to Cardiff University after application to the UK Biobank (project ref. 17044).

Based on our aim to investigate participants with no personal history of severe neuropsychiatric disorders, participants were removed if they were recorded as being affected by SZ, psychosis, autism spectrum disorder, dementia or intellectual disability across any of the following three diagnosis methods: self-reported diagnosis from a doctor at any assessment visit, self-reported diagnosis on the online follow-up mental health questionnaire, an ICD-10 hospital admission code for the relevant disorder; or if they self-reported other than white British and Irish descent. At this stage, 1,112 participants with neuropsychiatric disorders and 46,522 participants of non-white British and Irish descent were removed from the original sample (remaining n=454,985).

### Genotyping, CNV calling and CNV QC

Genotyping was performed using the Affymetrix UK BiLEVE Axiom array (807,411 probes) on an initial 50,000 participants, and the Affymetrix UK Biobank Axiom^®^ array (820,967 probes) on the rest of the sample. The two arrays have over 95% common content. Sample processing at UK Biobank is described in their documentation (https://biobank.ctsu.ox.ac.uk/crystal/docs/genotyping_sample_workflow.pdf).

CNV calling was conducted following the same procedure as described in a previous study^18^. Briefly, normalised signal intensity, genotype calls and confidences were generated using ~750,000 biallelic markers common to both arrays that were further processed with PennCNV-Affy software^20^. Individual samples were excluded if they had >30 CNVs, a waviness factor >0.03 or <−0.03 or call rate <96%. This quality control process resulted in the further exclusion of 13,652 individuals (3.0%; remaining n=441,333). Individual CNVs were excluded if they were covered by <10 probes or had a density coverage of less than one probe per 20,000 base pairs^18^.

### CNV annotation

To date, 12 CNVs have been significantly associated with SZ after correction for multiple testing^13–16^ and constituted the focus of this study (SZ-CNVs). A list of these 12 CNVs, and the genomic coordinates of their critical regions, can be found in supplemental table 1. The breakpoints of all SZ-CNVs included in our study were manually inspected to confirm that they met our CNV calling criteria (supplemental Table 1). Briefly, we required a CNV to cover more than half of the critical interval and to include the key genes in the region (if known), or in the case of single gene CNVs the deletions to intersect at least one exon and the duplications to cover the whole gene. As a control comparison, we used individuals that carried none of the 93 CNVs that have previously been associated with neurodevelopmental disorders^20,21^ (non-CNV carriers). The criteria for defining pathogenic CNVs has been previously fully described^18^.

### Brain imaging data

MRI data were collected in a single Siemens Skyra 3T scanner located at UK Biobank’s recruitment centre at Stockport (UK). Details of the brain imaging protocols have been published elsewhere^23^. Briefly, this project focuses on the 15 metrics of subcortical volumes - i.e. left and right thalamus, caudate, putamen, pallidum, hippocampus, amygdala and accumbens, overall ventricular volume – generated and provided by UK Biobank after applying FIRST (FMRIB’s Integrated Registration and Segmentation Tool^24^) to the T1-weighted brain images. At the outset of this study, brain MRI data were available for 9,112 individuals in our remaining sample (n=9,063 non-CNV carriers and n=49 SZ-CNVs carriers).

To avoid the potential effect of extreme values, outlier brain volumes – defined as values ±2.5 standard deviations from the group mean - in any subcortical brain structures were removed from the analyses. After removal of outliers, all brain measures were normally distributed and variance did not differ between groups. The volume of each subcortical structure was then z-transformed using the mean and standard deviation from the non-CNV carriers group as reference; therefore, individual z-scores for these variables represent the deviation in standard units from the non-CNV group’s mean.

### Cognitive test

During their MRI visit, participants underwent cognitive testing, including a measure of FI. The FI score corresponds to the total number of correct answers out of 13 verbal and numerical reasoning questions that participants were able to answer within 2 minutes. FI scores were available for 8,694 participants in our sample, were normally distributed and were also z-transformed in reference to the non-CNV group.

### Analyses

To investigate association between subcortical brain volumes and CNV-carrier status, regression analyses with the former as the predicted variable and the latter as the predictor were run for each brain structure. In an initial step, age, sex and brain size (total grey matter + white matter volume) were entered into the model, followed by CNV-carrier status. The significance threshold was set at a p<0.05 (two-sided), and False Discovery Rate (p.adjust in R^25^) was used to correct for multiple testing.

Brain volumes significantly associated with CNV-carrier status from the above analyses were taken into a mediation analyses, if they were also found to be significantly associated with performance in the FI test. Multiple mediation analysis was conducted using the latent variable analysis package (lavaan)^26^ for R. Age, sex and brain size were included as covariates and the covariation between mediators was also factored into the model.

## Results

Among SZ-CNVs carriers, 49% carried the most common 15q11.2deletion CNV. Our sample did not include carriers of the rarer SZ-CNVs (see table 1). Previous work from our group has shown consistency of individual CNV frequencies between UK Biobank batches, and between the UK Biobank and other independent control datasets^18^.

**Table 1.**
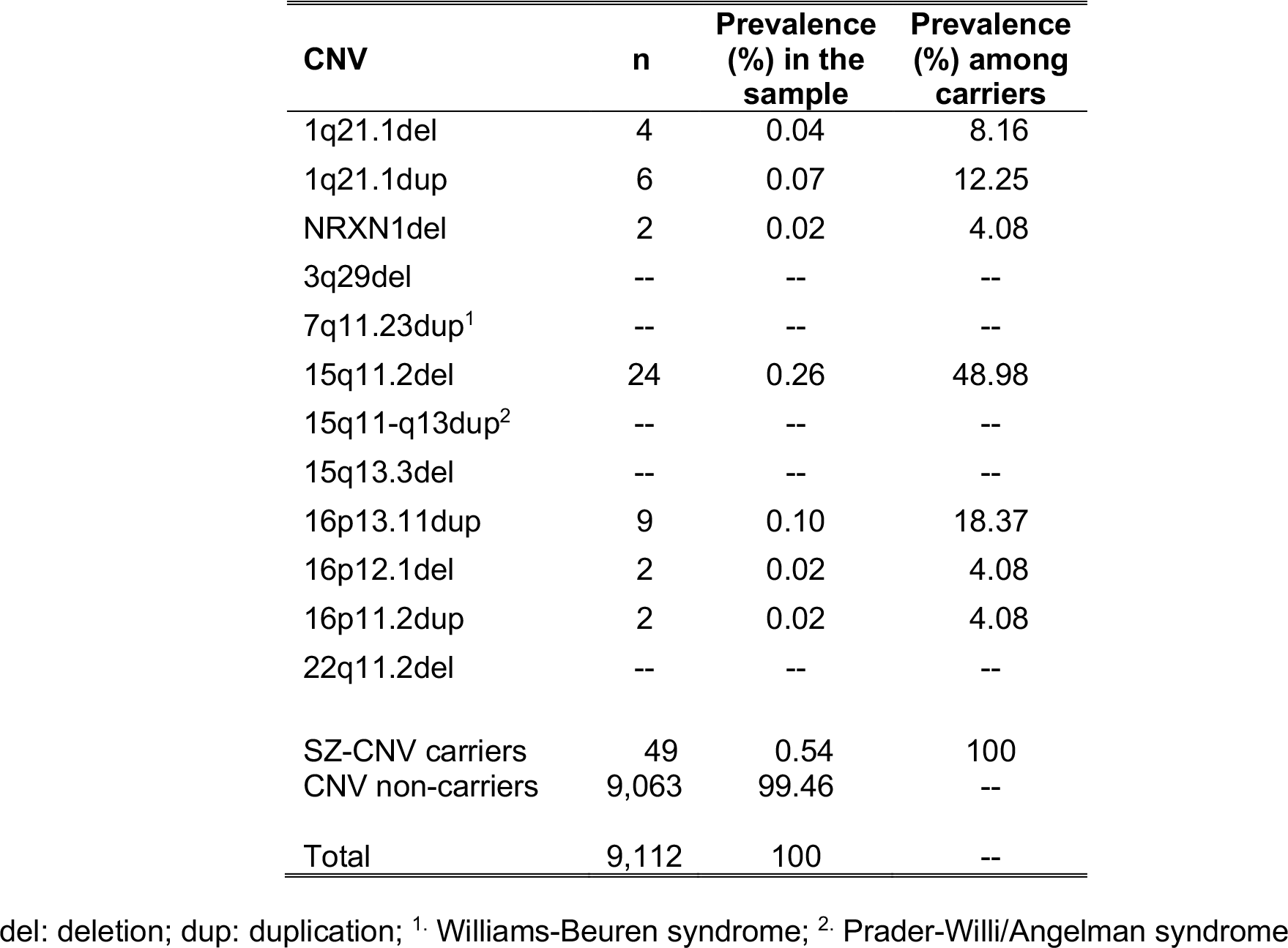
List of schizophrenia-associated copy number variants (CNVs) and their prevalence (%) in our sample

| **CNV** | **n** | **Prevalence (%) in the sample** | **Prevalence (%) among carriers** |
| --- | --- | --- | --- |
| 1q21.1del | 4 | 0.04 | 8.16 |
| 1q21.1dup | 6 | 0.07 | 12.25 |
| NRXN1del | 2 | 0.02 | 4.08 |
| 3q29del | -- | -- | -- |
| 7q11.23dup<sup>1</sup> | -- | -- | -- |
| 15q11.2del | 24 | 0.26 | 48.98 |
| 15q11-q13dup<sup>2</sup> | -- | -- | -- |
| 15q13.3del | -- | -- | -- |
| 16p13.11dup | 9 | 0.10 | 18.37 |
| 16p12.1del | 2 | 0.02 | 4.08 |
| 16p11.2dup | 2 | 0.02 | 4.08 |
| 22q11.2del | -- | -- | -- |
| SZ-CNV carriers | 49 | 0.54 | 100 |
| CNV non-carriers | 9,063 | 99.46 | -- |
| Total | 9,112 | 100 | -- |

del: deletion; dup: duplication; <sup>1</sup>. Williams-Beuren syndrome; <sup>2</sup>. Prader-Willi/Angelman syndrome

Participants’ mean age was 61.6 years (sd=7.04, range=45-73 years), with SZ-CNV carriers being approximately 2 years older than non-carriers (mean=63.5, sd=5.97 vs. mean=61.6, sd=7.04), t(48)=2.26, p=.029. Sex was equally distributed across group; Chi^2^(1)=0.68, p>.1; 52% female participants.

### Subcortical brain volumes association with SZ-CNVs carrier status

Five subcortical brain volumes showed significant association with SZ-CNV carrier status after correction for multiple testing (Table 2). In all these subcortical structures, carrying a SZ-CNV was predictive of a reduction in volume compared with non-CNV carriers. Only two subcortical structures showed no association with SZ-CNV carrier status in either left or right hemisphere: caudate and amygdala; and no differences were found either for bilateral ventricular volume (Table 2 and Figure 1). In order to ascertain whether these results were explained by the small age difference between groups, we selected a subgroup of non-CNV carriers that perfectly matched SZ-CNV carriers by age (t(48)=0.001, p=0.999) and the results remained the same (see supplemental table 2).

**Table 2.**
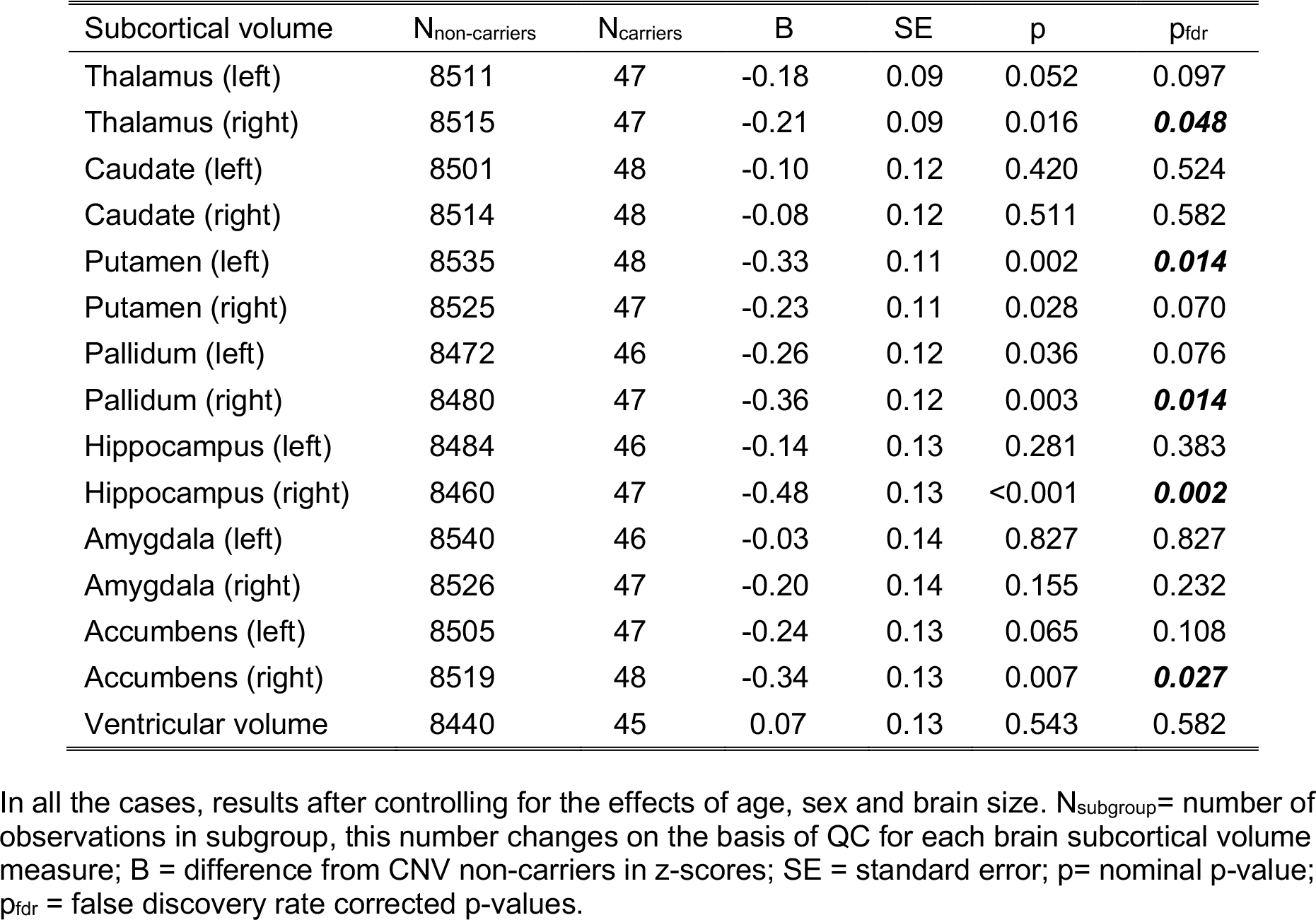
Summary statistics from linear regression analysis of the effect of schizophrenia-associated copy number variants (SZ-CNV) carrier status on the 15 subcortical brain volumes

**Figure 1.**
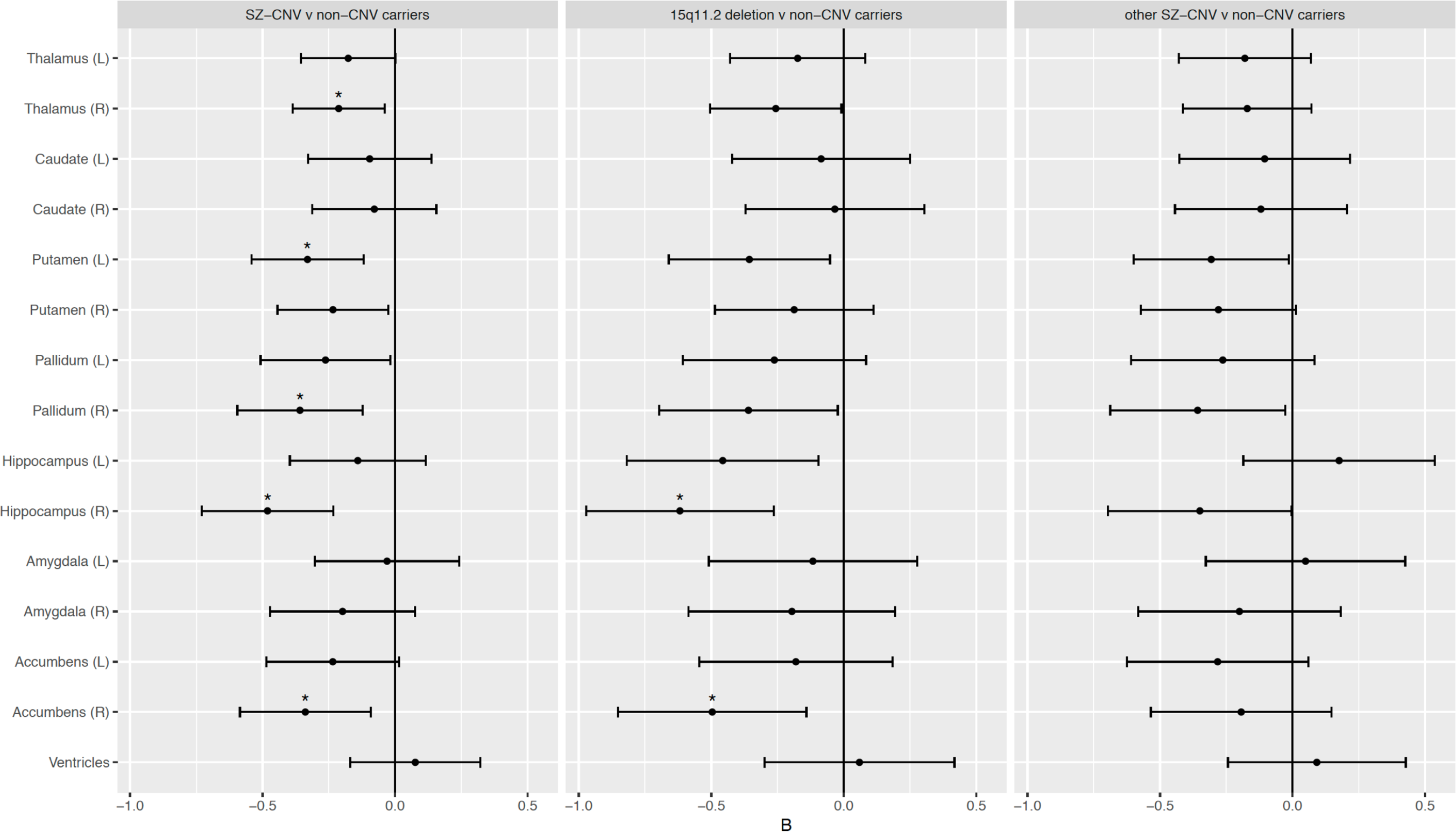
Differences in the volume of subcortical structures across SZ-CNV carriers v non-CNV carriers (left panel), 15q11.2deletion v non-CNV carriers (middle panel) and other SZ-CNV than 15q11.2deletion v non-CNV carriers (right panel). The x-axis reflects the standardised B value (z-score difference between groups, negative values indicate reduced volume in SZ-CNV carriers). Error bars indicate 95% confidence interval. Asterisks indicate differences that survive false discovery rate correction for multiple comparison.

To determine whether any of these effects were driven by the most frequent SZ-CNV in our sample – i.e. 15q11.2deletion – we tested subcortical brain volumes in 15q11.2deletion carriers vs. non-CNV carriers, and in carriers of any SZ-CNVs other than the 15q11.2deletion vs. non-CNV carriers. The results of these two tests replicated the above findings, with both CNV carrier groups showing the same direction of effects relative to non-carriers for all but one subcortical structure (Figure 1, supplemental tables 3 and 4). Left hippocampus, showed a nominally significant volume reduction in 15q11.2deletion carriers compared to non-CNV carriers (B=−0.46, SE=0.18, p=0.013, p_FDR_=0.066), whereas carriers of any other SZ-CNVs showed no significant difference (Figure 1, supplemental tables 3 and 4). A direct comparison between 15q11.2deletion carriers and carriers of any other SZ-CNVs showed hippocampal volume to be reduced in the former compared to the latter (B=−0.58, SE=0.27, p=0.035).

### Mediation of subcortical volume change over the association between SZ-CNV carrier status and cognitive performance

As expected from previous research^18^, SZ-CNV carriers performed worse in the FI test than non-CNV carriers (B=−0.55, SE=0.14, p<0.001). Moreover, all five brain volumes significantly associated with SZ-CNV carrier status were also significantly associated with FI score: right hippocampus, thalamus and accumbens, and left putamen and pallidum (Table 3). From the mediation analysis including these five volumes (Figure 2), there was a significant direct effect of SZ-CNV carrier status on FI (B=−0.46, p=0.002) and this association was partially mediated by the right thalamus (B=−0.04, p=0.009). The right hippocampus and left putamen showed a trend towards significance in mediating the association (B=−0.02, p=0.063; B=−0.02, p=0.060), whereas right accumbens and left pallidum were not significant. However, the total indirect effect of these five variables remained significant (B=−0.07, p<0.001), accounting for almost 14% (p=0.006) of the variance of the association between SZ-CNV carrier status and FI score.

**Table 3.** Summary statistics from linear regression analysis of the effect of the volume of subcortical structures associated with schizophrenia-associated copy number variants (SZ-CNV) carrier status on the performance (correct responses) in the fluid intelligence test.

| Subcortical volume | Cognitive test | n | B | SE | p |
| --- | --- | --- | --- | --- | --- |
| Thalamus (right) | Fluid<br>intelligence | 8202 | 0.09 | 0.02 | **<0.001** |
| Putamen (left) |  | 8225 | 0.04 | 0.01 | **0.004** |
| Pallidum (right) |  | 8167 | 0.03 | 0.01 | **0.032** |
| Hippocampus (right) |  | 8149 | 0.05 | 0.01 | **<0.001** |
| Accumbens (right) |  | 8202 | 0.02 | 0.01 | **0.050** |

In all the cases, results after controlling for age, sex and brain size.

**Figure 2.**
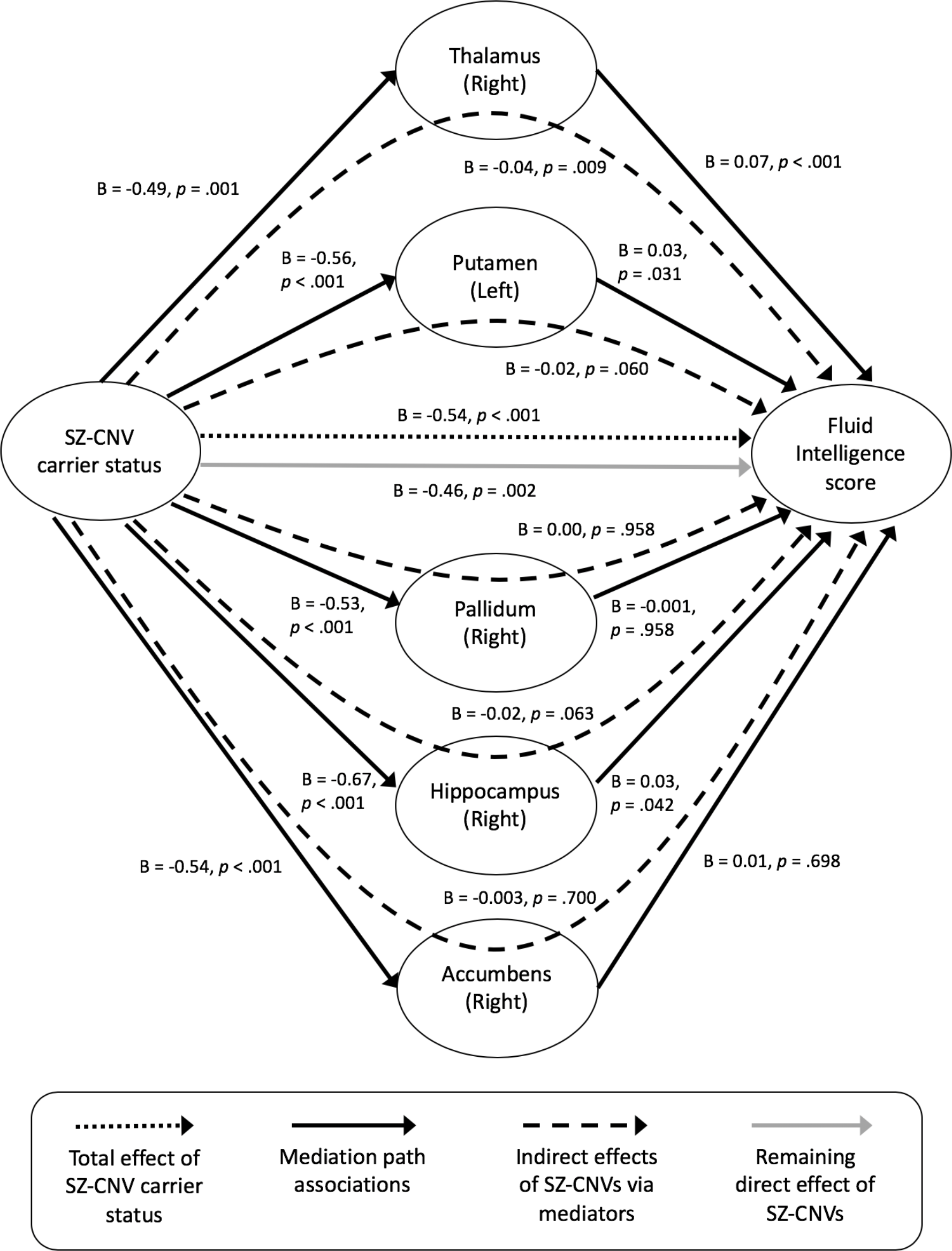
Results from the mediation analysis of brain volume over the association between SZ-CNV carrier status and performance in the fluid intelligence test. N= 7768 participants (44 SZ-CNV carriers v 7724 non-CNV carriers).

## Discussion

The main aim of this study was to ascertain whether unaffected SZ-CNV carriers present similar anatomical alterations in subcortical volumes to those previously found in clinical samples. Our results show this to be the case for thalamus, hippocampus and accumbens, whereas we find no association between SZ-CNV carrier status and amygdala, and an inverse association to that previously reported in clinical research with pallidum. Importantly, we show thalamic and hippocampal volumes to mediate the association between SZ-CNV carrier status and cognitive performance, demonstrating their relevance for explaining phenotypic variance.

Concordant with our hypothesis, we found evidence for a reduced volume in SZ-CNV carriers compared to non-CNV carriers in three subcortical brain structures previously found to be reduced in volume in SZ patients^7,8^, thus suggesting their potential to represent intermediate phenotypes for the disorder. Thalamus, accumbens and hippocampus in the right hemisphere were found to be smaller in SZ-CNV carriers after correction for multiple testing. Thalamus and accumbens in the left hemisphere showed a nominally significant reduction and a trend towards significance respectively, indicating the same direction of effect, albeit milder. Although the right hippocampus showed the strongest effect-size in our sample - almost half a standard deviation difference in volume between groups - the left hippocampus showed no association; suggesting clear interhemispheric differences that warrant further research. These results largely concur with recently reported results from two independent international consortia^7,8^, which showed the volumes of these three subcortical structures to be reduced in SZ patients compared to healthy controls. Moreover, subjects at high risk for SZ (i.e. first-degree relatives or prodromal individuals) have also shown reductions in these same structures compared to healthy samples^26–30^, with a recent meta-analysis also suggesting that hippocampal volumetric differences would be restricted to the right hemisphere^31^. Previous clinical research^7,8^ along with the results presented here suggest that the volume of the thalamus, the accumbens and the right hippocampus could represent genetically moderated premorbid risk markers for SZ, rather than reverse consequences of having the condition.

Contrary to our hypotheses, we found reduced volume in the pallidum in SZ-CNV carriers, and no group differences in amygdala or ventricular size. A previous study from our group^32^ showed a negative association between volume in the pallidum and risk for psychosis based on common variants in healthy volunteers, again opposing the results obtained from clinical samples. Like then, it can be argued that clinical samples can conceal the effects of illness related factors such use of antipsychotic drugs^33^, and this could explain the different results between clinical and at risk but otherwise healthy samples. In any case, our results indicate that previous volumetric differences observed in pallidum and amygdala between patients and controls are more likely to be a consequence of disorder status than a premorbid genetic marker of SZ. As per overall ventricular volume, the fact that lateral ventricular size was not available for this study – measure used in previous clinical studies - precludes any clear interpretation of our results. Future longitudinal studies following at risk populations into disorder development would be helpful in establishing the temporal sequence and potential causes of these anatomical changes.

Due to a sample size limitation, we were not able to study the potential effects of each individual SZ-CNVs, limiting our ability to provide insight into specific neurobiology associated with the disorder. However, we examined the differential effects of the most frequent SZ-CNV in our sample - 15q11.2deletion - against the rest of SZ-CNVs tested as a group. Overall, the pattern of results was very similar between 15q11.2deletion and the remaining SZ-CNVs, indicating that our overall group effects were not driven by this particular CNV. On the contrary, both carriers groups showed similar subcortical volumetric differences relative to non-CNV carriers, suggestive of a potential common pathobiological effect. In fact, supplementary analyses contrasting carriers of the 16p13.11 (n=9) and 1q21.1 (n=6) duplications against non-CNV carriers suggest similar effects - i.e. reductions in volume - in most subcortical structures, although no analysis for 16p13.11duplication reached significance (supplemental tables 5 and 6). Notably, carriers of the 15q11.2deletion showed a nominally significant reduction in left hippocampus volume compared to healthy controls, whereas carriers of any other SZ-CNVs showed no reduction. In a direct comparison between these two SZ-CNVs carrier groups, the volume of the left hippocampus was reduced in 15q11.2deletion carriers compared with any other SZ-CNVs carriers. This suggests that the left hippocampus could be one of the subcortical brain structures most susceptible to variation due to the causal mechanisms linked to rare genetic variation. If confirmed, this would help to inform a potential characterisation of phenotypic subtypes of SZ in a similar manner to a recent study which showed that hippocampal volume, taken bilaterally, characterises subtypes of SZ patients classified on the basis of cognitive decline^34^. However, the reduction in statistical power which resulted from dividing our sample into two CNV-carrier subgroups calls for caution when interpreting this result.

Significant associations between SZ genetic risk factors and brain biomarkers are of interest if they can translate into or modulate clinical phenotypes, since those have more potential to inform patient stratification and research into new treatment interventions. Previous research has shown impaired cognitive performance in SZ cases compared to control participants^17^, and in SZ-CNVs carriers^18,35,36^ compared to non-CNV carriers, this cognitive deficit (score in a FI test) also being present in our sample. A mediation analysis indicated that the volume of the five subcortical structures differing between SZ-CNV and non-CNV carriers explained a significant proportion of the association between SZ-CNV status and FI score, with the volume of the right thalamus showing the strongest effect. This result highlights the importance of investigating the impact of SZ-CNVs on brain anatomy, due to the potential for brain markers to mediate the effect of genetic risk on clinically relevant phenotypes. Our results do not discount that many other variables not examined in our study could also mediate this association; or that the right thalamus, or any of the other biomarkers identified here, could also mediate other CNV-phenotypic associations relevant for SZ, such as anhedonia. Since the SZ-CNVs considered here do not only increase risk for SZ but also for other neurodevelopmental disorders such as autism, it could be expected that any brain abnormalities associated with these CNVs would mediate the association between these rare variants and behavioural phenotypes relevant for these clinical diagnoses as a whole (as could be FI), following a cross-disorder perspective.

It is important to note that although we present here the results from the largest association study between CNVs and brain anatomy to date, the number of SZ-CNV carriers in our sample still limits the power of our analytical approaches. The fact that these genetic variants are rare resulted in fewer than 50 healthy carriers from an initial sample of ~10,000 participants. The UK Biobank project continues to acquire data and has set a target of ~100,000 scanned/genotyped participants to be completed within the next few years. This final sample will certainly improve our ability to explore genetic-biomarker-phenotypic associations, as well as providing replication samples for the results presented here. Also, despite the wealth of the UK Biobank data, phenotypic variables relevant for SZ or psychiatry in general are limited, which restricts its potential in this area of research. Finally, it is also important to note that the nature of recruitment in UK Biobank (volunteers put themselves forwards to be scanned) makes this resource not truly representative of the general population^37^, though it is difficult to see how this could result in type I error in the present study.

In summary, previous unsuccessful attempts to associate common genetic risk with subcortical brain volumes^11,38^ resulted in the hypothesis that these biomarkers could not associate with genetic risk for SZ. Here we prove that genetic risk factors associated with SZ, at least from the risk conferred by CNVs as opposed to common variants, are significantly associated with anatomical brain markers. Some of the associations found mirror previous results in clinical samples, highlighting the potential for these biomarkers to represent intermediate phenotypes for the disorder (i.e. volume in thalamus, accumbens and hippocampus); however, some of our results also oppose previous findings in clinical samples (i.e. pallidum and amygdala) indicating that these alterations are more likely to represent consequences, rather than premorbid markers, of the disorder. Importantly, we also show the above associations to mediate the correlation between genetic risk and a core clinical phenotype: cognitive performance.

## Acknowledgements

This research has been conducted using the UK Biobank Resource under Application Number 17044. KMK is supported by a Wellcome Trust Clinical Research Training Fellowship (ref. 201171/Z/16/Z). We thank Professor Sir Michael Owen for his comments on an early draft of this manuscript.

## Supplementary material

**Supplemental Table 1.** List of schizophrenia associated CNVs with critical region coordinates, calling criteria and number of genes hit. Schizophrenia (SZ) relative risk estimates for each CNV were taken from Rees *et al* 2016. PSD = Post-synaptic density, WBS = Williams-Beuren syndrome, PWS/AS = Prader-Willi/Angelman syndromes.

| Syndrome | Locus | Critical/Unique Sequence Region (hg19) | N Biallelic Axiom probes | PSD Genes (from 685 PSD genes) | Calling criteria | N genes | SZ relative risk |
| --- | --- | --- | --- | --- | --- | --- | --- |
| 1q21.1del | 1q21.1 | chr1:146,527,987-147,394,444 | 289 |  | Size >50% of critical region | 9 | 5.6 |
| 1q21.1dup | 1q21.1 | chr1:146,527,987-147,394,444 | 289 |  | Size >50% of critical region | 9 | 2.1 |
| *NRXN1*del | 2p16.3 | chr2:50145643-51259674 | 235 | *NRXN1* | Exonic deletions | 1 | 4.2 |
| 3q29del | 3q29 | chr3:195,720,167-197,354,826 | 627 | *DLG1* | Size >50% of critical region | 28 | 13.5 |
| WBSdup | 7q11.23 | chr7:72,744,915-74,142,892 | 349 | *STX1A* | Size >50% of critical region | 26 | 4.3 |
| 15q11.2del BP1-BP2 | 15q11.2 | chr15:22,805,313-23,094,530 | 145 | *CYFIP1* | Size >50% of critical region | 5 | 1.7 |
| PWS/ASdup | 15q11-q13 | chr15:22,805,313-28390339 | 1807 | *CYFIP1* | Full critical region, ~4Mbp | 116 | 29.2 |
| 15q13.3del BP4-BP5 | 15q13.3 | chr15:31,080,645-32,462,776 | 466 |  | Size >50% of critical region | 8 | 4.2 |
| 16p13.11dup | 16p13.11 | chr16:15,511,655-16,293,689 | 349 | *MYH11* | Size >50% of critical region | 7 | 1.7 |
| 16p12.1del (520kb) | 16p12.1 | chr16:21,950,135-22,431,889 | 168 | *UQCRC2* | Size >50% of critical region | 8 | 3.4 |
| 16p11.2dup (593kb) | 16p11.2 | chr16:29,650,840-30,200,773 | 193 | *ALDOA,CORO1A,MAPK3,TAOK2* | Size >50% of critical region | 30 | 8.6 |
| 22q11.2del | 22q11.2 | chr22:19,037,332-21,466,726 | 843 | *PI4KA,SEPT5* | Size >50% of critical region | 61 | 21.6 |

**Supplemental Table 2.**
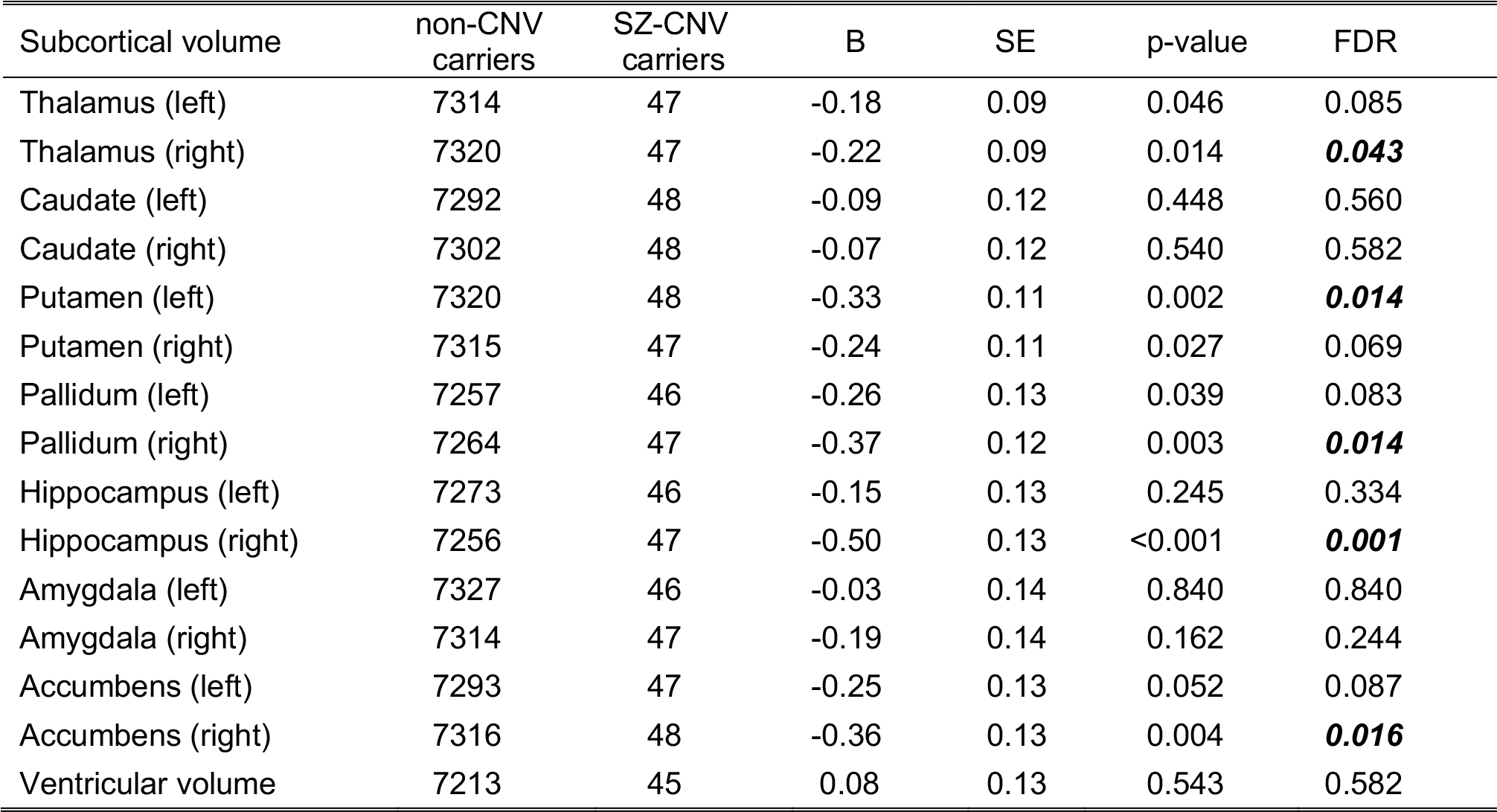
Summary statistics from linear regression analysis of the effect of SZ-CNV carrier status on the 15 subcortical brain volumes (subsample of non-carriers to carriers matched by age)

| Subcortical volume | non-CNV carriers | SZ-CNV carriers | B | SE | p-value | FDR |
| --- | --- | --- | --- | --- | --- | --- |
| Thalamus (left) | 7314 | 47 | -0.18 | 0.09 | 0.046 | 0.085 |
| Thalamus (right) | 7320 | 47 | -0.22 | 0.09 | 0.014 | **0.043** |
| Caudate (left) | 7292 | 48 | -0.09 | 0.12 | 0.448 | 0.560 |
| Caudate (right) | 7302 | 48 | -0.07 | 0.12 | 0.540 | 0.582 |
| Putamen (left) | 7320 | 48 | -0.33 | 0.11 | 0.002 | **0.014** |
| Putamen (right) | 7315 | 47 | -0.24 | 0.11 | 0.027 | 0.069 |
| Pallidum (left) | 7257 | 46 | -0.26 | 0.13 | 0.039 | 0.083 |
| Pallidum (right) | 7264 | 47 | -0.37 | 0.12 | 0.003 | **0.014** |
| Hippocampus (left) | 7273 | 46 | -0.15 | 0.13 | 0.245 | 0.334 |
| Hippocampus (right) | 7256 | 47 | -0.50 | 0.13 | <0.001 | **0.001** |
| Amygdala (left) | 7327 | 46 | -0.03 | 0.14 | 0.840 | 0.840 |
| Amygdala (right) | 7314 | 47 | -0.19 | 0.14 | 0.162 | 0.244 |
| Accumbens (left) | 7293 | 47 | -0.25 | 0.13 | 0.052 | 0.087 |
| Accumbens (right) | 7316 | 48 | -0.36 | 0.13 | 0.004 | **0.016** |
| Ventricular volume | 7213 | 45 | 0.08 | 0.13 | 0.543 | 0.582 |

**Supplemental Table 3.** Summary statistics from linear regression analysis of the effect of 15q11.2deletion carrier status on the 15 subcortical brain volumes

| Subcortical volume | non-CNV carriers | 15q11.2del carriers | B | SE | p-value | FDR |
| --- | --- | --- | --- | --- | --- | --- |
| Thalamus (left) | 8511 | 23 | -0.17 | 0.13 | 0.180 | 0.338 |
| Thalamus (right) | 8515 | 23 | -0.26 | 0.13 | 0.042 | 0.105 |
| Caudate (left) | 8501 | 23 | -0.09 | 0.17 | 0.617 | 0.712 |
| Caudate (right) | 8514 | 23 | -0.03 | 0.17 | 0.844 | 0.844 |
| Putamen (left) | 8535 | 23 | -0.36 | 0.16 | 0.022 | 0.082 |
| Putamen (right) | 8525 | 23 | -0.19 | 0.15 | 0.219 | 0.365 |
| Pallidum (left) | 8472 | 23 | -0.26 | 0.18 | 0.138 | 0.295 |
| Pallidum (right) | 8480 | 23 | -0.36 | 0.17 | 0.036 | 0.105 |
| Hippocampus (left) | 8484 | 23 | -0.46 | 0.18 | 0.013 | 0.066 |
| Hippocampus (right) | 8460 | 23 | -0.62 | 0.18 | 0.001 | **0.009** |
| Amygdala (left) | 8540 | 22 | -0.12 | 0.20 | 0.561 | 0.701 |
| Amygdala (right) | 8526 | 23 | -0.20 | 0.20 | 0.325 | 0.451 |
| Accumbens (left) | 8505 | 22 | -0.18 | 0.19 | 0.330 | 0.451 |
| Accumbens (right) | 8519 | 23 | -0.50 | 0.18 | 0.006 | **0.047** |
| Ventricular volume | 8440 | 21 | 0.06 | 0.18 | 0.747 | 0.800 |

**Supplemental Table 4.**
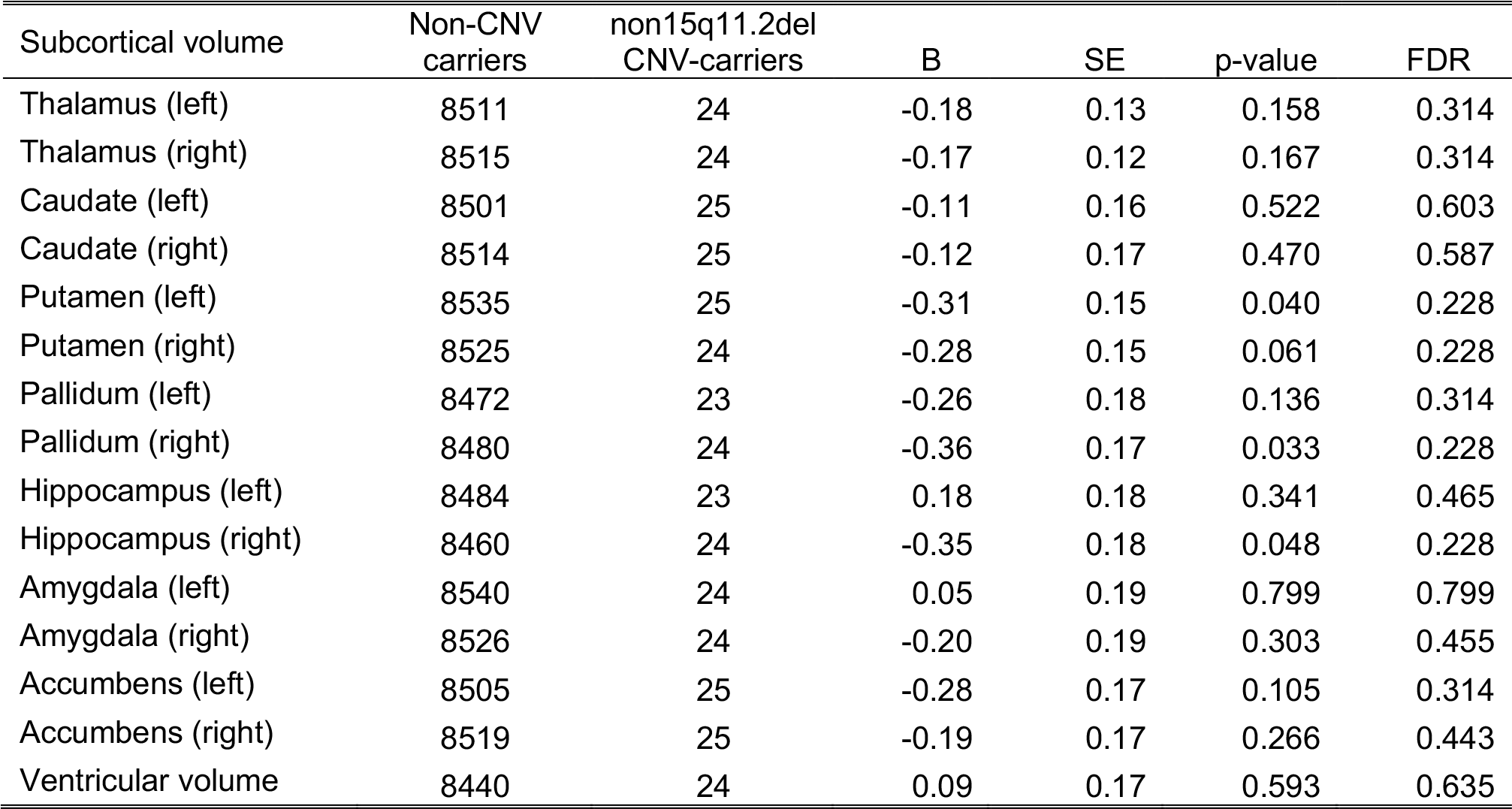
Summary statistics from linear regression analysis of the effect of carrying any SZ-CNV other than 15q11.2deletion status on the 15 subcortical brain volumes

| Subcortical volume | Non-CNV carriers | non15q11.2del CNV-carriers | B | SE | p-value | FDR |
| --- | --- | --- | --- | --- | --- | --- |
| Thalamus (left) | 8511 | 24 | -0.18 | 0.13 | 0.158 | 0.314 |
| Thalamus (right) | 8515 | 24 | -0.17 | 0.12 | 0.167 | 0.314 |
| Caudate (left) | 8501 | 25 | -0.11 | 0.16 | 0.522 | 0.603 |
| Caudate (right) | 8514 | 25 | -0.12 | 0.17 | 0.470 | 0.587 |
| Putamen (left) | 8535 | 25 | -0.31 | 0.15 | 0.040 | 0.228 |
| Putamen (right) | 8525 | 24 | -0.28 | 0.15 | 0.061 | 0.228 |
| Pallidum (left) | 8472 | 23 | -0.26 | 0.18 | 0.136 | 0.314 |
| Pallidum (right) | 8480 | 24 | -0.36 | 0.17 | 0.033 | 0.228 |
| Hippocampus (left) | 8484 | 23 | 0.18 | 0.18 | 0.341 | 0.465 |
| Hippocampus (right) | 8460 | 24 | -0.35 | 0.18 | 0.048 | 0.228 |
| Amygdala (left) | 8540 | 24 | 0.05 | 0.19 | 0.799 | 0.799 |
| Amygdala (right) | 8526 | 24 | -0.20 | 0.19 | 0.303 | 0.455 |
| Accumbens (left) | 8505 | 25 | -0.28 | 0.17 | 0.105 | 0.314 |
| Accumbens (right) | 8519 | 25 | -0.19 | 0.17 | 0.266 | 0.443 |
| Ventricular volume | 8440 | 24 | 0.09 | 0.17 | 0.593 | 0.635 |

**Supplemental Table 5.** Summary statistics from linear regression analysis of the effect of 16p13.11duplication carrier status on the 15 subcortical brain volumes

| Subcortical volume | Non-CNV carriers | 16p13.11dup CNV-carriers | B | SE | p-value | FDR |
| --- | --- | --- | --- | --- | --- | --- |
| Thalamus (left) | 8511 | 9 | 0.03 | 0.21 | 0.885 | 0.885 |
| Thalamus (right) | 8515 | 9 | 0.05 | 0.20 | 0.789 | 0.885 |
| Caudate (left) | 8501 | 9 | 0.33 | 0.27 | 0.233 | 0.672 |
| Caudate (right) | 8514 | 9 | 0.28 | 0.28 | 0.301 | 0.672 |
| Putamen (left) | 8535 | 9 | -0.14 | 0.25 | 0.562 | 0.844 |
| Putamen (right) | 8525 | 8 | -0.04 | 0.26 | 0.872 | 0.885 |
| Pallidum (left) | 8472 | 8 | -0.32 | 0.30 | 0.283 | 0.672 |
| Pallidum (right) | 8480 | 8 | -0.36 | 0.29 | 0.218 | 0.672 |
| Hippocampus (left) | 8484 | 9 | 0.53 | 0.29 | 0.073 | 0.672 |
| Hippocampus (right) | 8460 | 9 | -0.37 | 0.29 | 0.194 | 0.672 |
| Amygdala (left) | 8540 | 9 | 0.31 | 0.31 | 0.324 | 0.672 |
| Amygdala (right) | 8526 | 8 | -0.31 | 0.34 | 0.358 | 0.672 |
| Accumbens (left) | 8505 | 9 | -0.12 | 0.29 | 0.683 | 0.885 |
| Accumbens (right) | 8519 | 9 | -0.20 | 0.29 | 0.494 | 0.824 |
| Ventricular volume | 8440 | 8 | -0.06 | 0.30 | 0.838 | 0.885 |

**Supplemental Table 6.**
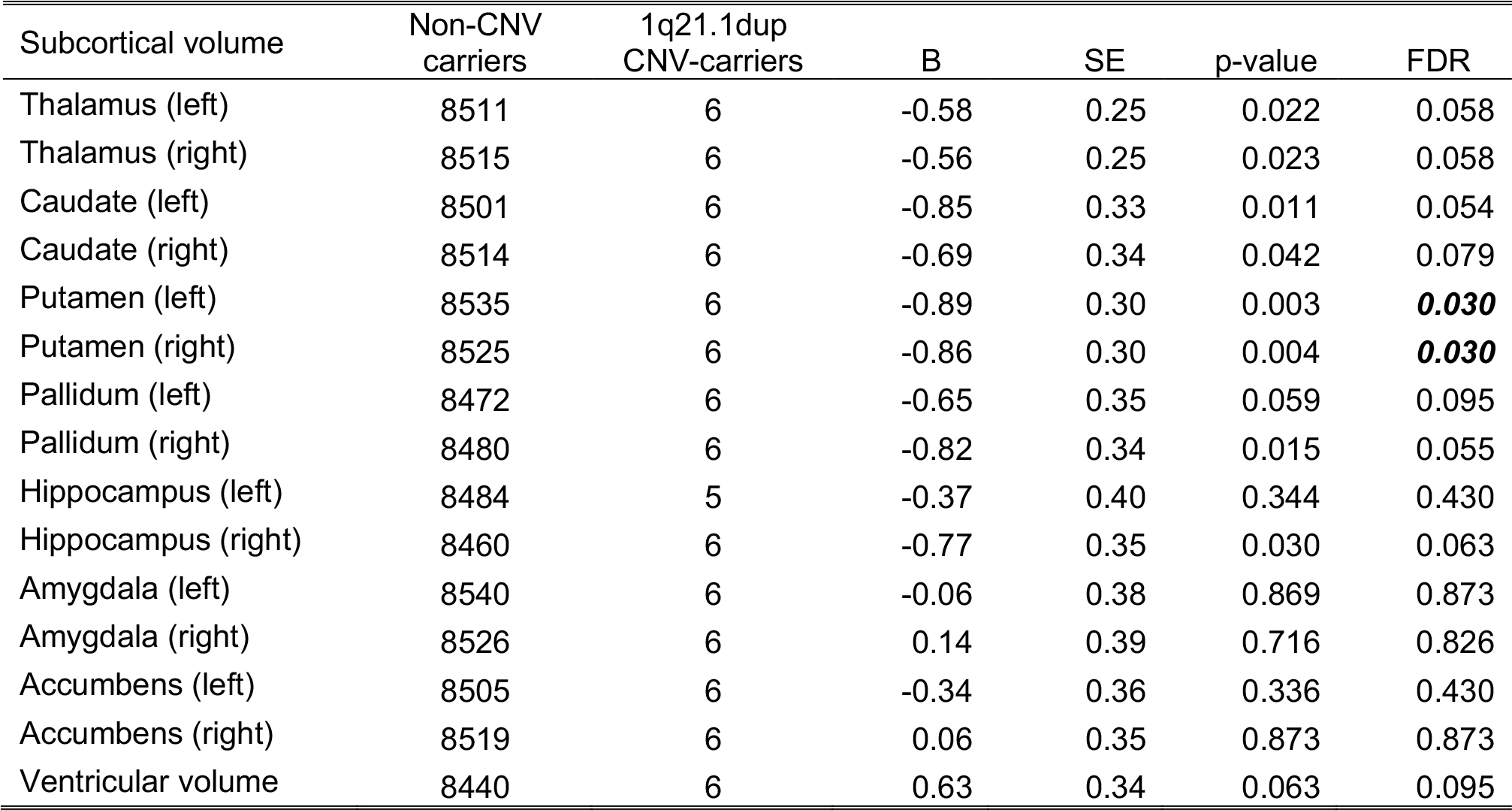
Summary statistics from linear regression analysis of the effect of 1q21.1duplication carrier status on the 15 subcortical brain volumes

| Subcortical volume | Non-CNV carriers | 1q21.1dup CNV-carriers | B | SE | p-value | FDR |
| --- | --- | --- | --- | --- | --- | --- |
| Thalamus (left) | 8511 | 6 | -0.58 | 0.25 | 0.022 | 0.058 |
| Thalamus (right) | 8515 | 6 | -0.56 | 0.25 | 0.023 | 0.058 |
| Caudate (left) | 8501 | 6 | -0.85 | 0.33 | 0.011 | 0.054 |
| Caudate (right) | 8514 | 6 | -0.69 | 0.34 | 0.042 | 0.079 |
| Putamen (left) | 8535 | 6 | -0.89 | 0.30 | 0.003 | **0.030** |
| Putamen (right) | 8525 | 6 | -0.86 | 0.30 | 0.004 | **0.030** |
| Pallidum (left) | 8472 | 6 | -0.65 | 0.35 | 0.059 | 0.095 |
| Pallidum (right) | 8480 | 6 | -0.82 | 0.34 | 0.015 | 0.055 |
| Hippocampus (left) | 8484 | 5 | -0.37 | 0.40 | 0.344 | 0.430 |
| Hippocampus (right) | 8460 | 6 | -0.77 | 0.35 | 0.030 | 0.063 |
| Amygdala (left) | 8540 | 6 | -0.06 | 0.38 | 0.869 | 0.873 |
| Amygdala (right) | 8526 | 6 | 0.14 | 0.39 | 0.716 | 0.826 |
| Accumbens (left) | 8505 | 6 | -0.34 | 0.36 | 0.336 | 0.430 |
| Accumbens (right) | 8519 | 6 | 0.06 | 0.35 | 0.873 | 0.873 |
| Ventricular volume | 8440 | 6 | 0.63 | 0.34 | 0.063 | 0.095 |

